# RNA editing and trans-splicing with reprogrammed tracrRNAs

**DOI:** 10.64898/2026.06.28.735058

**Authors:** Adini Q. Arifah, Constantinos Patinios, Rachael Larose, Chase L. Beisel

**Author notes:** Correspondence (to C.L.B.).

## Abstract

Natural CRISPR-Cas9 systems rely on crRNA-tracrRNA duplexes to guide DNA targeting. Prior work showed that tracrRNAs could be reprogrammed to hybridize to cellular RNAs, resulting in their conversion into non-canonical crRNAs that enabled RNA detection and recording. However, the fate of the cellular RNA and the engineering opportunities it affords remain unexplored. Here, we show that the hybridized RNA is not inactivated, allowing the recruitment of Cas9 to the RNA duplex to drive RNA base editing and trans-splicing. Fusing ADAR2dd to dSpyCas9 and systematically engineering the reprogrammed tracrRNA (Rptr) enabled efficient and tunable A-to-I RNA editing, with on- and off-target profiles comparable to dCas13. The methodology extended to the compact CjeCas9 that could be further tailored for RNA targeting by deleting the HNH domain and mutating the PAM-interacting domain. Finally, utilizing Rptrs to block splicing enabled 3′ and 5′ RNA trans-splicing. Thus, Rptrs offer a versatile alternative to conventional Cas9 guide RNA architectures for programmable RNA manipulation.

## INTRODUCTION

In Type II CRISPR–Cas systems, guide RNA activity depends on a tracrRNA that hybridizes with the crRNA, together forming the functional RNA complex required for targeting by Cas9. Beyond this canonical role, tracrRNAs can also hybridize with RNAs containing repeat-like sequences, converting them into non-canonical CRISPR RNAs (ncrRNAs) that recruit Cas9 for DNA targeting^1^. Building on this principle, reprogrammed tracrRNAs (Rptrs) have been developed to hybridize RNAs of interest as ncrRNAs and have enabled a range of technologies, including RNA detection (LEOPARD^1^, PUMA^2^ and AGATHA^3^) and RNA recording (TIGER^4^ and CHEETAH^5^).

Within these advances, Rptr-based approaches have so far converted target RNAs into crRNA-like molecules that direct Cas9 or related nucleases toward DNA substrates^1–5^. It thus remains unknown whether a Rptr–target RNA complex can instead recruit Cas9 to introduce functional changes on the captured target RNA itself. Although, naturally RNA-targeting CRISPR–Cas systems such as Cas13 have enabled programmable RNA cleavage and editing^6–10^, Cas9 remains an attractive platform for programmable RNA targeting because it is extensively characterized, highly engineerable, and available in diverse orthologs. Previous attempts to repurpose Cas9 for RNA targeting have consistently relied on sgRNA, including strategies using PAMmer oligonucleotides, modified guide RNAs, or engineered Cas9 variants^11–13^. What remains unexplored is whether the RNA-targeting capability of Rptrs could be utilized for this same purpose.

Here, we establish a programmable RNA engineering platform based on Rptr-guided recruitment of catalytically inactive Cas9 to RNA substrates. By fusing dCas9 to the ADAR2 deaminase domain and guiding it with Rptrs designed to hybridize to target RNAs, we achieve site-specific Adenosine-to-Inosine (A-to-I) RNA editing in human cells. This strategy is compatible with both SpyCas9 and CjeCas9 and does not require removal of DNA-targeting domains, though in some cases such deletions support protein size reduction with minimal effect on editing activity. Beyond RNA base editing, we further demonstrate that Rptr-guided dCas9 enables programmable RNA trans-splicing, allowing the introduction of large sequence changes into target transcripts. Together, these results establish Rptrs as a versatile platform for RNA engineering and extend their functionality beyond RNA detection and recording toward direct manipulation of cellular RNAs.

## RESULTS

### Rptrs enable Cas9–mediated RNA editing

To determine whether Rptrs can direct Cas9 to edit the hybridized RNA, we evaluated Rptrs ability to recruit the catalytically inactive *Streptococcus pyogenes* Cas9 (dSpyCas9) fused at the C-terminus with the hyperactive mutant (E488Q) deaminase domain of human ADAR2 (ADAR2dd) (**Fig. 1A**)^14^. ADAR2 is an adenosine deaminase acting on RNA, converting adenosine to inosine in double-stranded RNA (dsRNA) molecules. The fusion protein was separated by a nuclear export signal (NES) to mediate nuclear export of the fusion protein and a GSGGGGS linker (dSpyCas9–ADAR2dd). As a positive control, we used REPAIR-v1, consisting of nuclease-dead PspCas13b fused to ADAR2dd (dPspCas13b–ADAR2dd) (**Fig. 1B**)^15^. Rptrs were designed based on the original *S. pyogenes* Cas9 tracrRNA architecture used for DNA targeting. Each Rptr was 38 nucleotides (nt) long, retained the canonical 5′-AAGU-3′ bulge sequence, and contained an adenosine–cytidine (A–C) mismatch opposite the target adenosine to promote its deamination by ADAR2dd^1^ (**Fig. 1C**). To restrict activity only to RNA when using dSpyCas9, we selected targets of which corresponding genomic DNA sites lacked a 5′-NGG-3′ PAM (immediately 3′ of the protospacer on the non-target strand), thereby preventing Cas9 binding to DNA which might reduce Cas9 availability for RNA targeting (**Fig. 1C**). We refer to this system as ***Re****programmed **t**racrRNA for **R**NA **E**ngineering **A**pplica**t**ions using **Sp**yCas9* (RETREAT-Sp).

**Fig. 1.**
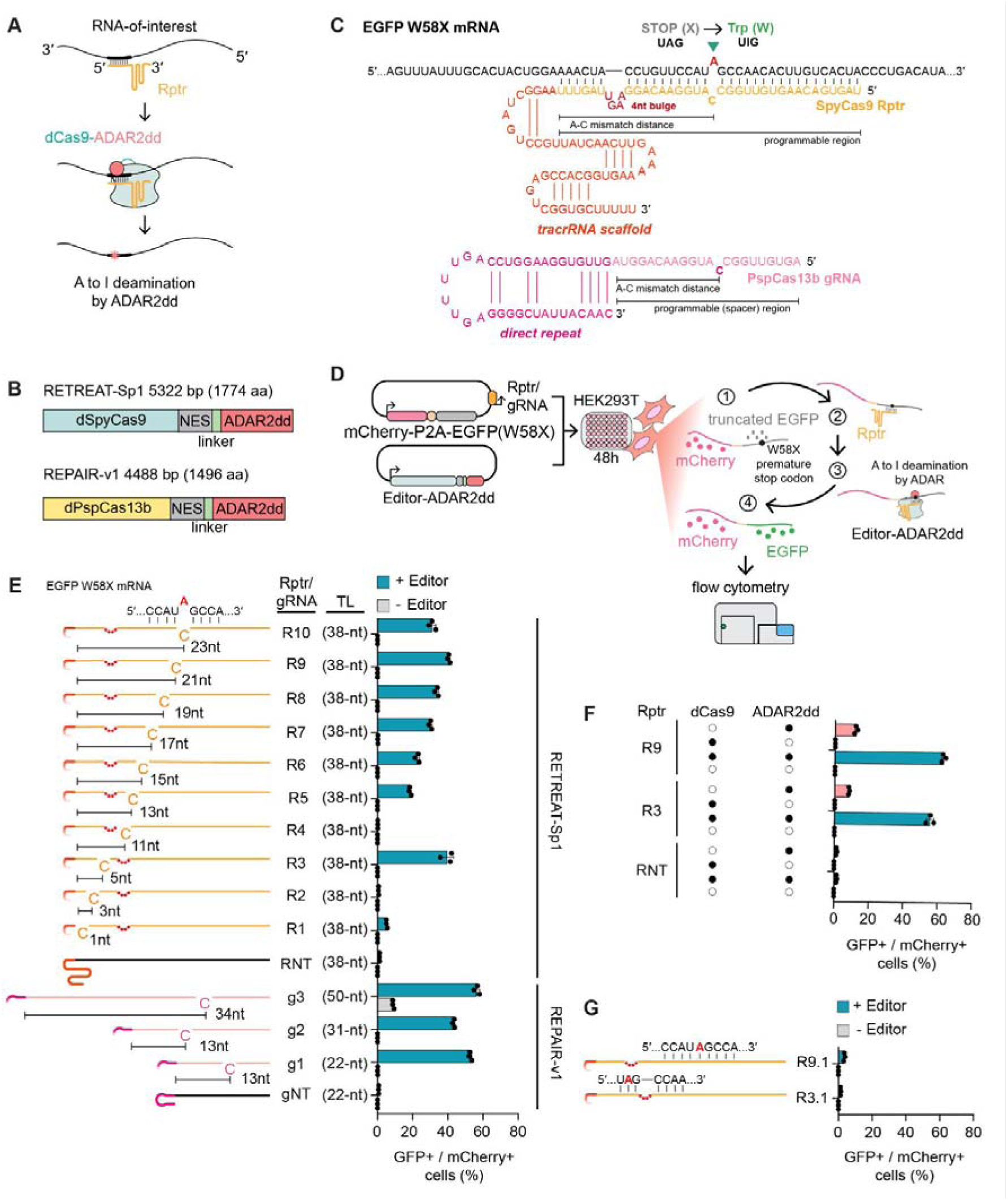
RETREAT enables programmable RNA base editing. (**A**) RETREAT concept: the reprogrammed tracrRNAs (Rptrs) hybridize to the target RNA and recruit dCas9–ADAR2dd for A-to-I editing. (**B**) Domain organization of the dSpyCas9–ADAR2dd (RETREAT-Sp1) and dPspCas13b–ADAR2dd (REPAIR-v1) editors. NES, nuclear export signal; linker, GSGGGGS. (**C**) Example design of a SpyCas9 Rptr targeting EGFP(W58X), in which the programmable region replaces the anti-repeat and the downstream scaffold enables Cas9 loading (top). Representative PspCas13b gRNA targeting EGFP(W58X) (bottom). The A–C mismatch distance is measured from the 3′ end of the programmable region for both Rptrs and gRNAs. (**D**) Workflow of the EGFP rescue assay. The mCherry–P2A–EGFP(W58X) reporter bearing the Rptr or gRNA is co-transfected with the respective editor. Editing converts UAG to UIG, restoring EGFP, detected by flow cytometry. (**E**) Assessment of the protein components required for EGFP restoration. Colored dots indicate the presence and open dots the absence of each component. Bars show the percentage of EGFP-positive cells with ADAR2dd alone (pink), dSpyCas9 alone (gray), or the dSpyCas9–ADAR2dd fusion (teal). (**F**) Assessment of different protein components for EGFP rescue. Colored dots indicate presence and empty dots absence of components. Bars show percentage of EGFP-positive cells for ADAR2dd only (pink), dSpyCas9 only (gray) or the fusion (teal). (**G**) Removal of the A–C mismatch abolishes EGFP rescue. Bars show the percentage of EGFP-positive cells in the presence (teal) or absence (gray) of the editor. For E, F and G, data represent mean ± SD (n = 3), with individual replicates shown as dots.

To assess RNA editing activity, we used an enhanced green fluorescent protein (EGFP) reporter in HEK293T cells in which the W58 codon (UGG) was replaced with a premature stop codon (UAG) (EGFP W58X) that abolishes fluorescence. Rptrs were designed to base pair with the EGFP mRNA at this position such that the adenosine in the UAG stop codon was opposite a cytosine in the Rptr, creating an A-C mismatch. This mismatch promotes deamination by ADAR2dd, converting adenosine to inosine, which is interpreted as guanosine during translation. The resulting UIG codon restores EGFP fluorescence, which we quantified by flow cytometry (**Fig. 1D**). To measure transfection efficiency, an mCherry reporter was placed upstream of EGFP W58X and separated by a P2A sequence, allowing EGFP rescue to be quantified within the mCherry-positive population by flow cytometry (**Fig. S1, Supplementary Dataset 1**).

For initial evaluation, ten Rptr designs were tested in RETREAT-Sp1 against the EGFP-W58X reporter, varying the position of the A-C mismatch relative to the start of the tracrRNA scaffold **(Fig. 1E**). Except for Rptr 2 (R2) and Rptr 4 (R4), all designs increased EGFP rescue relative to both the non-targeting control and the no-editor condition, confirming RETREAT-Sp1–dependent activity. Among the active guides, Rptr 3 (R3) and Rptr 9 (R9), which place the A-C mismatch at positions 5 and 21, respectively, produced the highest percentages of EGFP-positive cells (∼40%). For comparison, the dPspCas13b–ADAR2dd editor produced ∼53% and ∼56% EGFP-positive cells with the 22-nt and 50-nt spacers (g1 and g3), respectively (**Fig. 1E**). EGFP restoration was abolished in the absence of the fused ADAR2dd-Cas9 or the A–C mismatch and was strongly reduced when ADAR2dd was expressed separately, indicating that reporter rescue depends on both local recruitment of ADAR2dd-Cas9 and the designed mismatch opposite the target adenosine (Fig. 1F,G). Together, these results validate RETREAT-Sp1 as a functional RNA base editing platform and establish initial design principles, especially the positioning of the A-C mismatch.

### RNA base editing is influenced by the Rptr architecture

We next asked whether additional modifications to the Rptr design could influence EGFP reporter activation. As a first step, we altered the overall length of the Rptr and measured EGFP-positive cells using the same flow cytometry assay. Shortening the 5′ end of Rptr 3 to 28-nt reduced EGFP rescue from 41% to 26% (R3.2) (**Fig. S2**). The same truncation had a stronger effect on Rptr 9, reducing EGFP rescue from 43% to 19% (R9.2), potentially because its A-C mismatch is positioned closer to the 5′ end of the Rptr (**Fig. S2**). Unexpectedly, extending the Rptr length to 53-nt and 63-nt also impaired activity, reducing EGFP rescue by Rptr 3 from 41% to 35% (R3.4) and 36% (R3.6), and Rptr 9 from 43% to 24% (R9.4) and 30% (R9.6). These extensions also increased editor-independent activity, with the 63-nt variants producing 5% and 9% EGFP-positive cells in the absence of the editor for Rptr 3 and Rptr 9, respectively (**Fig. 2A**–**B**).

**Fig. 2.**
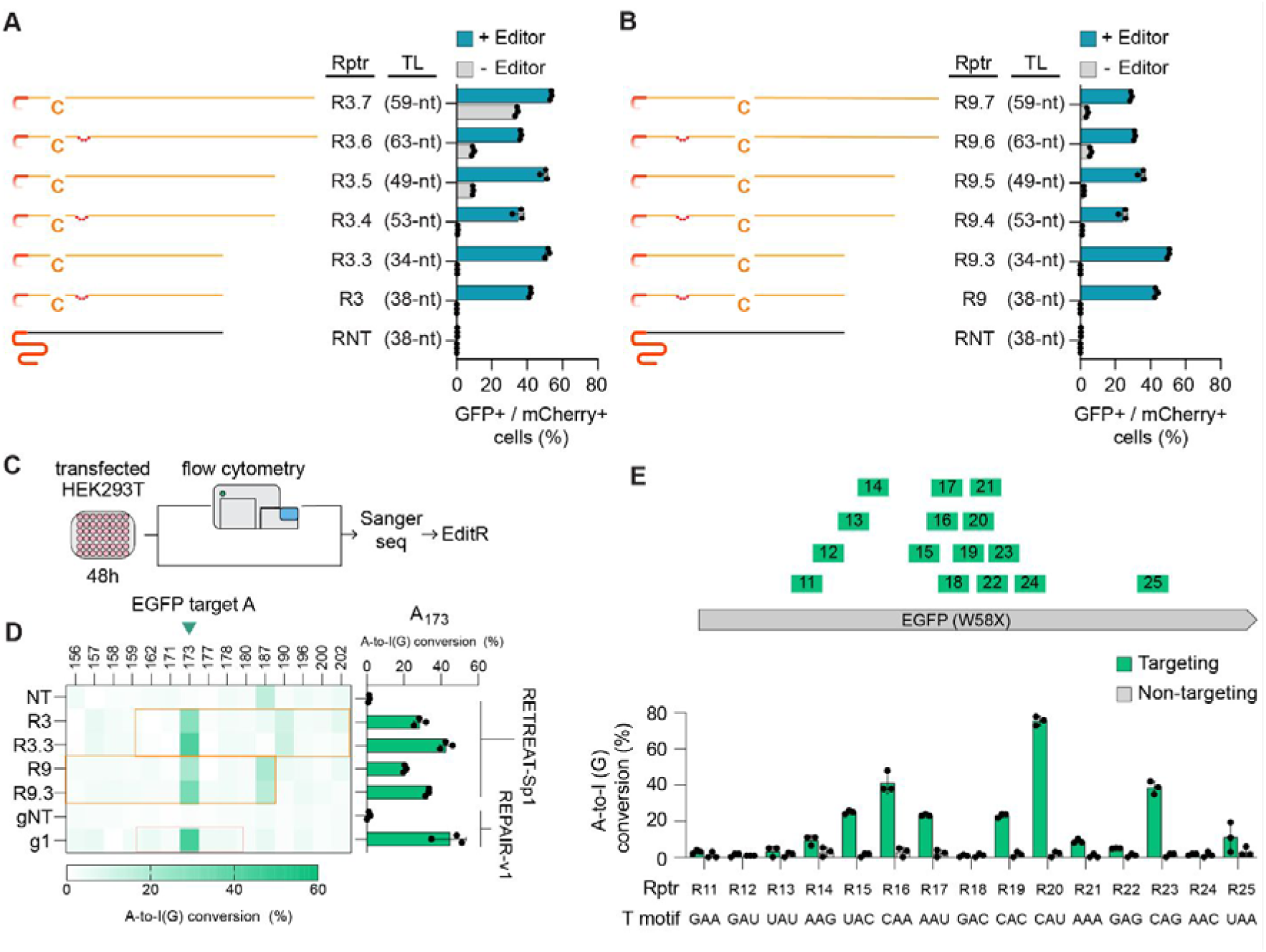
Bulge-free Rptrs enhance editing by RETREAT and allow multi-site targeting. (**A**) Flow cytometry analysis of Rptr 3 variants and (**B**) Rptr 9 variants. Rptrs have varying 5′ lengths with or without the 5′-AAGU-3′ bulge. Rptr design is shown on the left of each bar plot. Bars show the mean percentage of EGFP-positive cells with (teal) or without (gray) the editor. (**C**) Workflow for RNA editing analysis. EGFP rescue was assessed by flow cytometry. Samples were also analyzed by Sanger sequencing to obtain editing profiles. (**D**) A-to-I(G) conversion % as defined through Sanger sequencing. Adenosines proximal to the EGFP(W58X) site are shown, with boxed regions indicating complementarity to Rptrs. Heatmaps show mean A-to-I(G) conversion % (n = 3); bar graphs show the mean A-to-I(G) conversion of the target A173. (**E**) Base editing along the EGFP(W58X) target RNA. Target sites are shown at the top. Editing efficiencies (A-to-I(G)) quantified by Sanger sequencing and EditR (y-axis) are plotted for each Rptr (x-axis), alongside the corresponding 5′-NAN-3′ triplet motif flanking the edited adenosine, labeled in the figure as the *T motif* (bottom). For A, B, D and E, bar plots represent mean ± SD (n = 3), with individual replicates shown as dots.

We next investigated the role of the 5′-AAGU-3′ bulge, a structural element of the canonical *S. pyogenes* tracrRNA required for DNA cleavage^16,17^. Because the basis of this requirement remains unclear, the bulge could be dispensable for RNA editing or could instead be required to form a functional Rptr–Cas9 complex. We therefore tested whether removing it could simplify the Rptr architecture without compromising activity. To test this, we deleted the 5′-AAGU-3′ bulge from the original Rptr 3 and Rptr 9, resulting in Rptr 3.3 and Rptr 9.3. Bulge removal increased EGFP rescue by ∼10% for both designs (**Fig. 2A–B**).

Interestingly, the combined effect of bulge removal and 5′ extensions differed between the two designs. For Rptr 3.3, adding 15 nucleotides (R3.5) or 25 nucleotides (R3.7) markedly increased editor-independent EGFP rescue from 0.3% to 9% and 34%, respectively, while yielding no editor-dependent activity beyond the bulge-deleted version of the initial length (R3.3). In comparison, for Rptr 9.3, the 15-nt (R9.5) and 25-nt (R9.7) extensions reduced EGFP rescue from 50% to 35% and 29%, respectively, while background EGFP rescue remained low. (**Fig. 2A–B**).

Sanger sequencing confirmed that the bulge-deleted variants increased A-to-I(G) conversion by approximately 12–14%. For Rptr 9 (with bulge) and Rptr 9.3 (bulge-deleted), the predominant bystander edit occurred at A187, similar to what was observed with the non-targeting Rptr. In contrast, Rptr 3 (with bulge) and Rptr 3.3 (bulge-deleted) produced a distinct bystander profile, with A190 emerging as the major edited site (**Fig. 2C-D**).

Finally, to test whether RETREAT-Sp1 could edit additional sites within the EGFP-W58X transcript, we evaluated Rptr 11 – Rptr 25 which were based on variant 9.3 (bulge-deleted, A-C mismatch positioned 17 nt from the scaffold), targeting adenosines at 15 different sites. Editing outcomes were quantified by Sanger sequencing, showing a variation in editing efficiency across target sites. Approximately 50% of the target positions showed detectable activity (above the 10% Sanger threshold^18^) and most sites contained <45% the desired A-to-I(G) conversion (**Fig. 2E**). Together, these findings show that Rptr architecture strongly influences RETREAT-Sp1 activity, with bulge removal improving editing without increasing background, whereas changes in Rptr length generally reduced activity and increased editor-independent editing.

### RETREAT-Sp1 can edit endogenous RNAs with on- and off-target effects comparable to existing technologies

To evaluate endogenous RNA editing, we developed a single-plasmid system encoding both the editor and Rptr and targeted three endogenous transcripts: the abundant housekeeping gene ACTB, the naturally occurring ADAR2 target CYFIP2, and the therapeutically relevant oncogene KRAS^12,19^. All were expressed in HEK293T cells, while the selected adenosines showed little or no basal editing, allowing targeted editing to be measured with minimal background.

Rptrs were based on the EGFP Rptr 9 scaffold and designed with or without the bulge sequence. A-to-I(G) editing was quantified by targeted amplicon sequencing and normalized to transfection efficiency, which was determined from EGFP fluorescence. EGFP was fused to the C-terminus of the editor and separated through a self-cleaving P2A peptide (**Figs. 3A and S3A**).

**Fig. 3.**
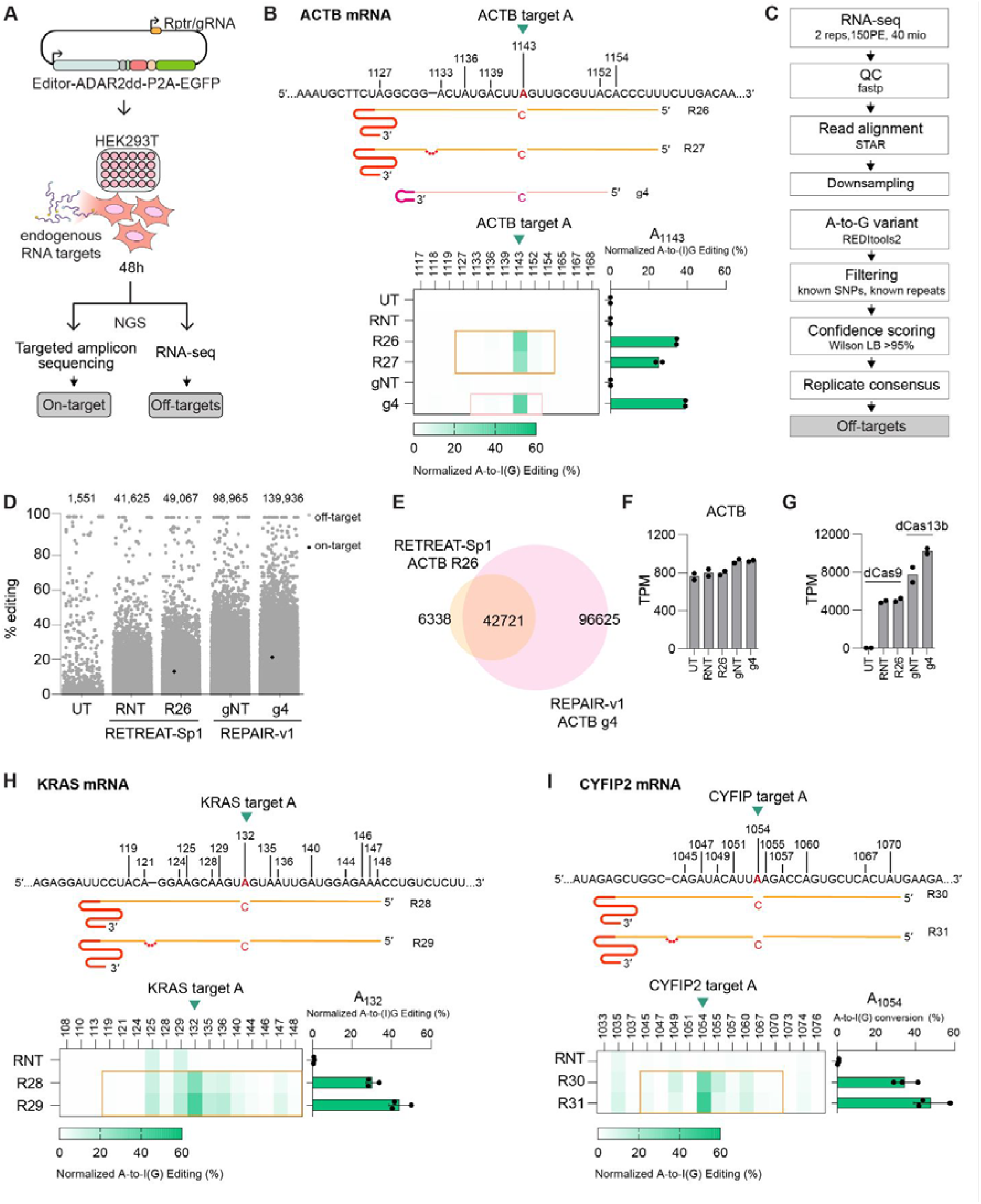
RETREAT edits endogenous transcripts with bystander edits and off-target profiles comparable to REPAIR-v1. (**A**) Schematic of the single-plasmid RNA base editing system. HEK293T cells were transfected with a plasmid containing the editor (dSpyCas9/dPspCas13b)–ADAR2dd–P2A–EGFP and the respective guide RNA (Rptr/gRNA), followed by NGS of amplicons to quantify on-target and off-target editing. (**B**) Editing of endogenous ACTB RNA normalized to transfection efficiency. Heatmaps show the mean A-to-I(G) editing efficiency at adenosines surrounding the target site, while bar graphs show editing at the target adenosine, A1143. (**C**) Schematic of computational workflow for transcriptome-wide off-target identification. (**D**) Transcriptome-wide A-to-I(G) edits shown as a dot plot in untransfected cells and in cells transfected with REPAIR-v1 or RETREAT-Sp1 using targeting or non-targeting guides. (**E**) Venn diagram showing unique and shared off-target sites for REPAIR-v1 with ACTB targeting gRNA 4 and RETREAT-Sp1 with ACTB targeting Rptr 26. (**F**) Abundance of ACTB mRNA in untransfected (UT) cells and in cells transfected with REPAIR-v1 using targeting (g4) or non-targeting (gNT) guides or RETREAT-Sp1 using targeting (R26) or non-targeting (RNT) guides, shown as transcripts per million (TPM). (**G**) Abundance of dPspCas13b–ADAR2dd and dSpyCas9–ADAR2dd transcripts following transfection with targeting or non-targeting guides, shown as TPM. (**H**) Editing of the endogenous KRAS RNA normalized to transfection efficiency. Heatmaps show the mean A-to-I(G) editing efficiency at adenosines surrounding the target site; bar graphs show editing of the target adenosine, A132. (**I**) Editing of the endogenous CYFIP2 RNA normalized to transfection efficiency. Heatmaps show the mean A-to-I(G) editing efficiency at adenosines surrounding the target site; bar graphs show editing of the target adenosine, A1054. For B, F, and G, bar plots represent mean (n =2), with individual replicates shown as dots. For H and I, bar plots represent mean ± SD (n = 3), with individual replicates shown as dots.

For ACTB, RETREAT-Sp1 with the bulge-free Rptr 26 (R26) achieved 35% normalized editing at A1143, compared to 24% for the bulged Rptr 27 (R27), consistent with the trend observed in the EGFP reporter assay (**Fig. 3B**). REPAIR-v1 achieved a comparable normalized editing efficiency of 39% (**Fig. 3B**), indicating that RETREAT-Sp1 and REPAIR-v1 have similar editing capacities at this target. Low levels of bystander editing were observed for both RETREAT-Sp1 and REPAIR-v1 systems targeting ACTB (**Fig. 3B**).

Since off-target activity is a well-known effect of RNA editing systems such as REPAIR-v1^12,15,20^, we next assessed the global specificity of RETREAT-Sp1. REPAIR-v1 was included as a benchmark, which includes the original PspCas13b-based RNA editor before subsequent efforts to improve specificity at the expense of efficiency^15^. Off-target profiles of REPAIR-v1 using gRNA 4 (g4) and RETREAT-Sp1 using the bulge-free Rptr (R26), both targeting the same region at the ACTB mRNA, were analyzed by performing directional RNA-seq. Following, transcriptome-wide off-target identification was performed following the computation workflow illustrated in **figure 3C**.

This analysis identified 1,551 off-targets in untransfected control; 98,965 and 139,936 off-targets in cells transfected with REPAIR-v1 using a non-targeting (gNT) or targeting gRNA (g4), respectively; and 49,067 and 41,625 off-targets in cells transfected with RETREAT-Sp1 using a non-targeting (RNT) or targeting Rptr (R26), respectively (**Fig. 3D**). Of the off-target sites detected for RETREAT-Sp1 using R26, 87% (42,721) were also present in REPAIR-v1 using g4 whilst 13% (6,338) were unique to RETREAT-Sp1 (**Fig. 3E, Dataset S2**). Around 1.3% were shared between the untransfected control and RETREAT-Sp1 using R26, and 60.6% were shared between RNT and R26 (**Fig. S4**). In contrast, only 31% (42,721) of the off-targets in REPAIR-v1 using g4 were shared with RETREAT-Sp1 using R26, and the majority of the edits (96,625) from REPAIR-v1 were unique. Overall, REPAIR-v1 exhibited approximately 2.9-fold and 2.4-fold more off-target sites than RETREAT-Sp1 under targeting and non-targeting conditions, respectively.

Because RNA targeting could affect target transcript stability, we examined ACTB abundance. ACTB expression remained similar across RETREAT-Sp1, REPAIR-v1, and the corresponding control conditions, indicating that RETREAT-Sp1 did not measurably reduce target RNA levels even under targeting conditions (**Fig. 3F**). We then compared editor abundance to assess whether differences in expression could contribute to the observed editing profiles. We showed that dPspCas13b-ADAR2dd was expressed at approximately two-fold higher levels than dSpyCas9-ADAR2dd (TPM ≈ 5,000 vs. 10,000) (**Fig. 3G**). The higher editor abundance may partly explain the greater number of off-target sites detected with REPAIR-v1 using both targeting and non-targeting guides. Taken together, these results show that RETREAT-Sp1 produces comparable on-target editing and lower off-target editing levels compared to REPAIR-v1 under the tested conditions.

Lastly, to examine whether the on-target editing trend observed for ACTB extends to other endogenous transcripts, we targeted KRAS and CYFIP2 for editing. Both genes are expressed at low levels in HEK293T cells, with ACTB expression approximately 52-fold and 84-fold higher than that of KRAS and CYFIP2, respectively (**Fig. S3b**). Both KRAS and CYFIP2 mRNA targets showed higher on-target editing with the bulged Rptrs (R29 for KRAS and R31 for CYFIP2) compared to the bulge-free variants (R28 and R30), yielding 44% versus 30% for KRAS and 48% versus 34% for CYFIP2 transcripts, respectively (**Fig. 3H–I**). KRAS mRNA exhibited substantial bystander editing, likely due to its high adenosine density within the Rptr-binding region, whereas CYFIP2 showed fewer bystander edits (**Fig. 3H-I**). Thus, RETREAT-Sp1 can edit various endogenous transcripts.

### Rptr-guided RNA editing is preserved across Cas9 orthologs and minimized Cas9 variants

After establishing RETREAT-mediated RNA editing with dSpyCas9, we sought to generate a more compact editor by testing whether domains required for DNA targeting could be removed. We focused on *Campylobacter jejuni* Cas9 (CjeCas9), a naturally compact Cas9 ortholog whose tracrRNA-derived Rptr is only 25 nt long and lacks a bulge (**Fig. 4A,B**) ^1^. Full-length dCjeCas9 (RETREAT-Cj1) supported Rptr-mediated RNA editing, with the efficiency of EGFP rescue depending on the position of the A–C mismatch within the Rptr–target RNA duplex (**Fig. 4C**).

**Fig. 4.**
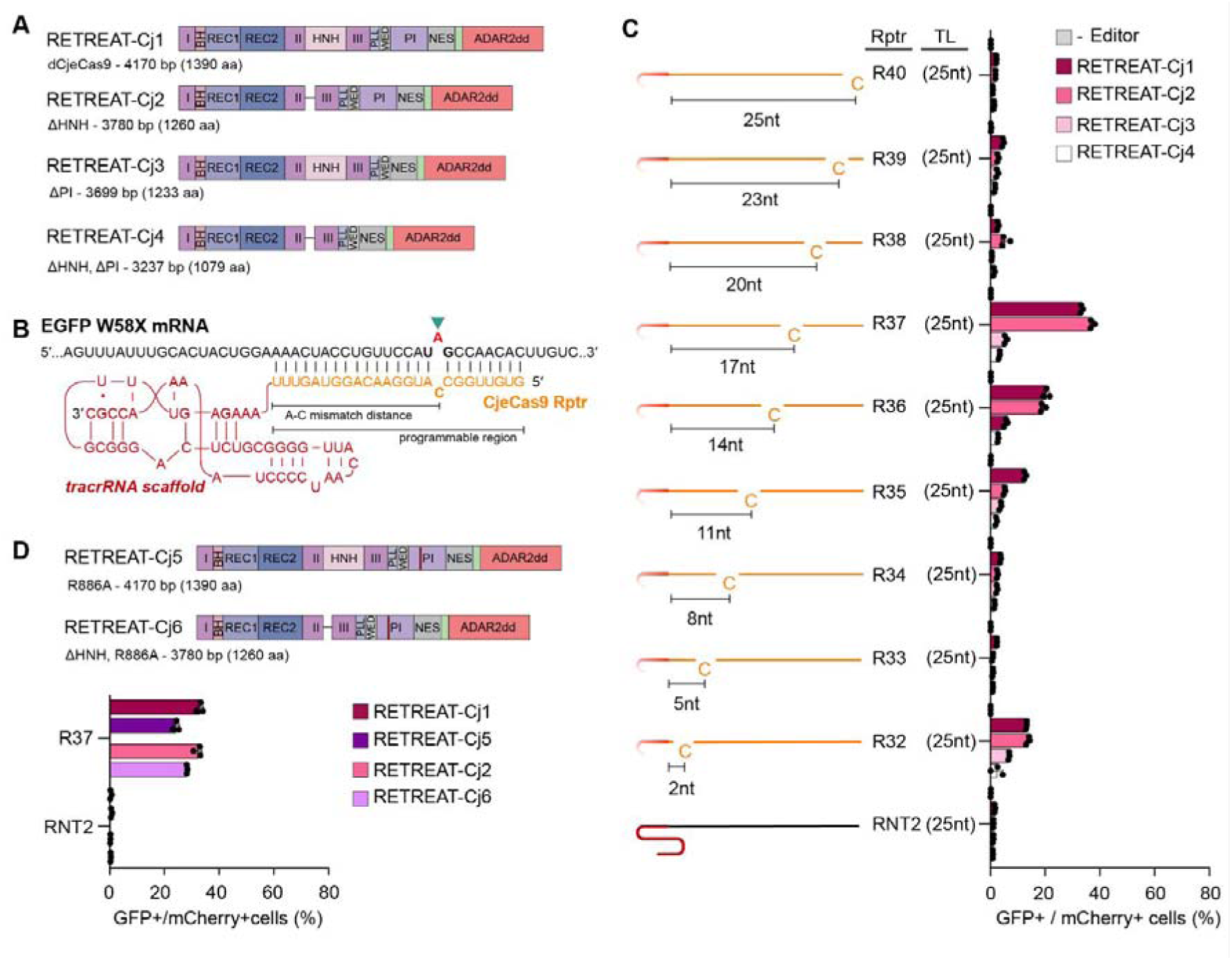
RNA editing directed by Rptrs is preserved among Cas9 orthologs and engineered variants. (**A**) Schematic of CjeCas9-based editors: RETREAT-Cj1 (full length), RETREAT-Cj2 (ΔHNH), RETREAT-Cj3 (ΔPI), RETREAT-Cj4 (ΔHNH/ΔPI), with GGGSGG linkers at deletions. (**B**) Representative CjeCas9 Rptr targeting the EGFP(W58X) site. (**C**) EGFP rescue by RETREAT-Cj1–Cj4 and a no-editor control, tested with Rptr 32–40 and RNT2. Bars show the mean percentage of EGFP-positive cells. (**D**) Schematic and EGFP rescue of PI domain mutants RETREAT-Cj5 (R866A) and RETREAT-Cj6 (ΔHNH, R866A) compared with RETREAT-Cj1 and RETREAT-Cj2 using R37 and a non-targeting control. Bars show the mean percentage of EGFP-positive cells. For C and D, bar plots represent mean ± SD (n = 3), with individual replicates shown as dots.

Given that the HNH domain mediates DNA cleavage but is not expected to contribute to RNA editing, we replaced it with a linker previously shown to preserve Cas9 structure^21^, generating RETREAT-Cj2. This alteration reduced the editor coding sequence by 360 bp and increased EGFP rescue using Rptr 37 (R37) from 28.9% to 37.2%, demonstrating that the HNH domain is dispensable for RETREAT-mediated RNA editing.

We next asked whether the PAM-interacting (PI) domain could also be removed to further reduce editor size, as PAM recognition is not required for RNA targeting. However, deletion of the PI domain, either alone or together with HNH deletion, reduced EGFP rescue to near-background levels (**Fig. 4C**). In contrast, mutation of R866, a PI-domain residue involved in PAM recognition and efficient DNA cleavage ^21^ reduced but did not abolish activity, decreasing EGFP rescue from 33.0% to 24.2% in full-length dCjeCas9 and from 32.2% to 28.2% in the ΔHNH variant when using R37 (**Fig. 4D**). Similarly, deletion of the SpyCas9 PI domain strongly impaired editing. In contrast, the R1333A and R1335A substitutions, previously shown to disrupt PAM recognition and DNA cleavage^22^, were largely tolerated and produced Rptr-dependent effects (**Fig. S5**), suggesting that the structural integrity of the PI domain is more important for RNA editing than PAM recognition by individual residues. These findings establish CjeCas9 as a compact RETREAT system, identify HNH deletion as a strategy to further reduce editor size without compromising activity, and indicate that PI domain integrity is more important for RNA editing than PAM recognition by individual residues.

### Rptr-guided dSpyCas9 mediates 3′ and 5′ RNA trans-splicing

Deaminase-based editors are limited to single-nucleotide transitions, restricting their utility for applications that require larger RNA sequence changes^23,24^. Inspired by recent studies using CRISPR-associated proteins for RNA repair and splice modulation^25–29^, we then asked whether RETREAT could be extended beyond RNA base editing to facilitate RNA trans-splicing.

We first established a 3′ trans-splicing split-EGFP reporter assay in HEK293T cells, in which successful trans-splicing restores EGFP expression (**Fig. 5A–B**). In this system, a trans-splicing molecule (TM) containing a binding domain (BD) and a minimal hemi-intron was recruited to the target RNA through RNA hybridization. Transfected cells were identified using a two-dimensional gate on the mCherry and BFP fluorescence plot to isolate the double-positive population. Trans-splicing efficiency was quantified as the percentage of GFP-positive cells within this mCherry+/BFP+ gate (**Fig. S6**). Targeting Rptrs alone modestly increased trans-splicing across all tested BDs compared with non-targeting controls, suggesting that Rptr–RNA interactions can influence splice outcomes independently of dSpyCas9 (**Fig. 5B**). Recruitment of dSpyCas9 enhanced trans-splicing for all three BDs. BD2 showed the largest absolute increase, from approximately 9% to 16% EGFP-positive cells, whereas BD3 reached the highest overall level, increasing from approximately 16% to 22% (**Fig. 5C**). Amplicon sequencing confirmed that targeting Rptrs increased trans-splicing, with larger effects in the presence of dSpyCas9. The efficiency increased from approximately 1% to 2% for BD2 and from 6% to 9% for BD3 without dSpyCas9, compared with 6% to 18% and 7% to 25%, respectively, with dSpyCas9. Although effect sizes differed from the fluorescence readout, both assays showed preferential enhancement with targeting Rptrs and identified BD3 as the most efficient design (**Fig. 5C**). Rptr design was guided by prior optimization of the TM and Rptr architectures, which showed that Rptr configuration affects trans-splicing efficiency (**Fig. S7**).

**Fig. 5.**
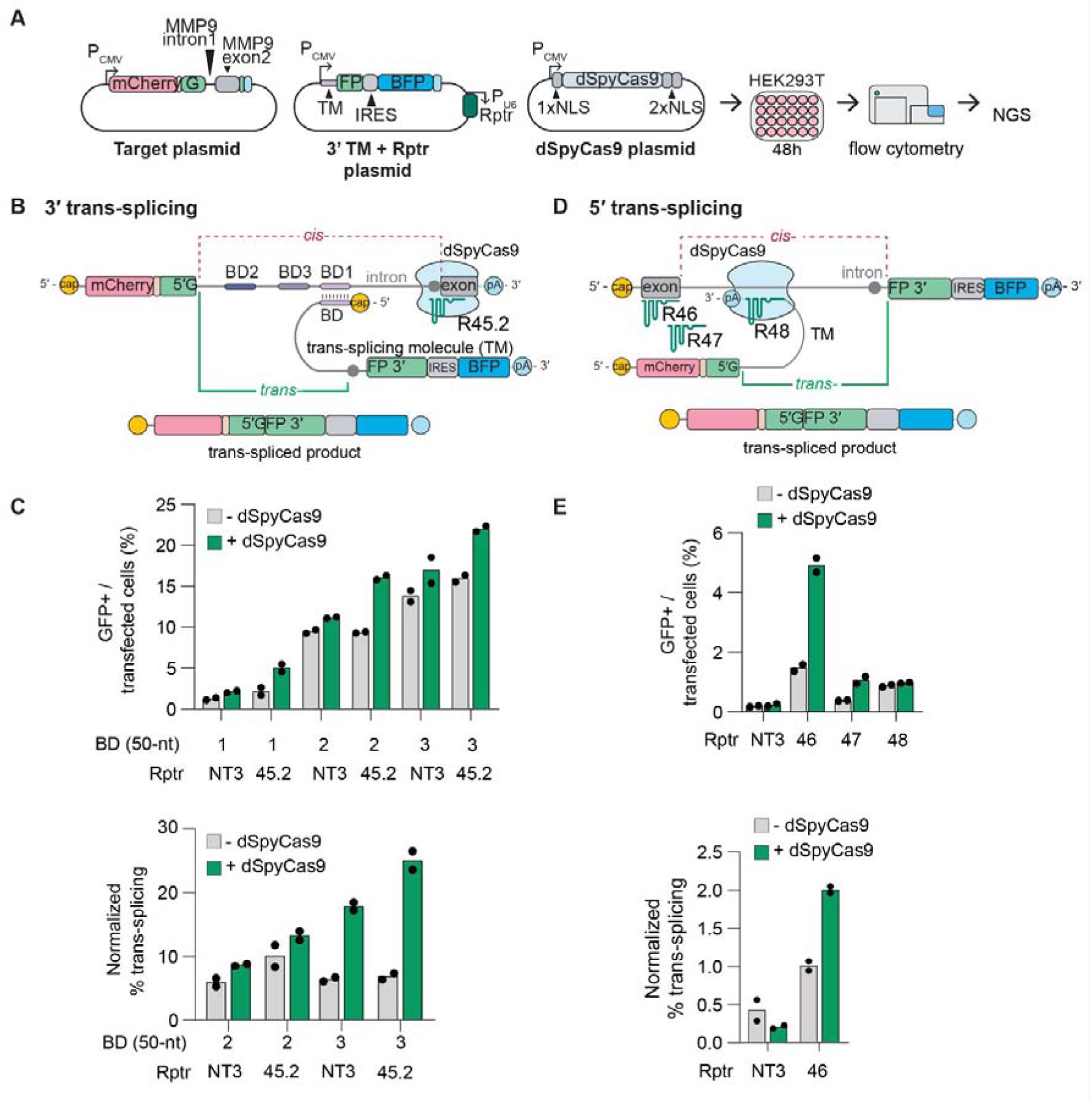
Rptr-guided dSpyCas9 facilitates 3′ and 5′ RNA trans-splicing. (**A**) Illustration of plasmids and methods used for 3′ trans-splicing. Plasmids carrying the target reporter (donor), the Rptr + TM and 3′ cargo (acceptor), and the dCas9-NLS, were transformed to HEK293T cells and subjected to flow cytometry and NGS. (**B**) Schematic of 3′ trans-splicing using extended binding domains (BD1–BD3, 50 nt) with Rptr 45.2. Controls include a non-targeting Rptr and omission of dCas9. (**C**) Flow cytometry analysis showing GFP-positive cells within the population positive for both mCherry and BFP, indicating trans-splicing efficiency (top). NGS-based quantification of trans-spliced products, shown as the percentage of trans-spliced reads relative to the sum of trans-spliced and cis-spliced products, normalized to transfection efficiency (percentage of cells positive for both mCherry and BFP among live cells) (bottom). (**D**) Schematic of 5′ trans-splicing with Rptr fused to the hemi-intron and cargo. Rptrs 46–48 and non-targeting controls were tested. (**E**) Flow cytometry analysis showing GFP-positive cells within the population positive for both mCherry and BFP, indicating trans-splicing efficiency (top). NGS-based quantification of trans-spliced products, shown as the percentage of trans-spliced reads relative to the sum of trans-spliced and cis-spliced products, normalized to transfection efficiency (percentage of cells positive for both mCherry and BFP among live cells) (bottom). For B and C, bar plots represent mean (n = 2), with individual replicates shown as dots.

We next examined whether RETREAT could also support 5′ trans-splicing using an analogous split-EGFP reporter system. Initial designs similar to 3’ trans-splicin with separate TM and Rptr showed no effect with RETREAT (**Fig. S8A-B**). However, fusing Rptr 46 directly to the TM produced a dSpyCas9-dependent increase in trans-splicing (**Fig. 5D**). EGFP-positive cells increased from approximately 1.5% without dSpyCas9 to 4.9% with dSpyCas9, compared with approximately 0.2% for the non-targeting control under both conditions (**Fig. 5E**). Amplicon sequencing confirmed the targeting-dependent effect in the presence of dSpyCas9, with trans-splicing increasing from approximately 1.0% for the non-targeting control to 2.0% for Rptr 46 (**Fig. 5E**). In this design, we used TM variants containing BD3, which provided the highest basal trans-splicing signal (**Fig. S9**). Together, these results establish that RETREAT can be extended beyond RNA base editing to support programmable RNA trans-splicing.

## DISCUSSION

Targeted RNA engineering can correct pathogenic transcripts without permanently altering the genome^15,30^. Several CRISPR systems, including Cas13, Cas9, Cas7-11, IscB variants, and type III-A complexes, have been adapted for RNA knockdown, base editing, and splicing^12,13,15,29,31^. While many approaches rely on naturally RNA-targeting effectors^6–9^, others repurpose Cas9 using PAMmers or engineered guide RNAs and protein domains to enable RNA targeting and reduce DNA activity^11–13^. A common theme in these examples is the reliance on the guide RNA for target RNA recognition. Our work pursued an entirely distinct approach by relying on reprogrammed tracrRNAs to achieve RNA editing and rewriting.

Using dCas9–ADAR2dd, RETREAT enabled efficient A-to-I editing of reporter and endogenous transcripts in human HEK293T cells without measurably altering target-transcript expression. Editing efficiency was strongly influenced by the position of the A–C mismatch within the Rptr–target duplex. Across both SpyCas9 and CjeCas9 systems, centrally positioned mismatches were generally favored, whereas scaffold-proximal mismatches were poorly edited. This pattern is consistent with structural occlusion of proximal bases within the Cas9-bound guide region, which would reduce access of ADAR2dd to the edited adenosine^17,21^. Editing also declined when the mismatch was moved toward the distal terminus, suggesting that duplex stability at the guide 5′ end is also important. Together, this dataset defines a positional window in which target accessibility and RNA hybrid stability are jointly optimized.

When exploring the architecture of the Rptr, one surprising finding was the impact of removing the 5′-AAGU-3′ bulge. Previous work showed that converting the repeat–anti-repeat bulge of the sgRNA into a perfectly base-paired duplex abolished DNA cleavage *in vitro* as well as indel formation in human cells^16^. As crystal structures implied that the bulge was part of how SpyCas9 recognizes the crRNA-tracrRNA duplex^17^, removing the bulge was expected to also strongly impair Rptr-mediated RNA editing. However, we found the opposite, with removing the bulge retaining or even enhancing the editing efficiency (**Figs. 2A–B, 3B**). This finding suggests that the bulge plays a role downstream of Cas9 binding the crRNA-tracrRNA duplex, such as part of DNA binding or cleavage. There is also the possibility that the lack of a bulge reduces the affinity of Cas9 for the duplex, which lends to transient recruitment of the dCas9-ADAR fusion, adenine deamination, and then release.

This target dependence extended beyond bulge removal. Across multiple EGFP sites and endogenous targets, editing efficiencies varied substantially and were not explained solely by transcript abundance, indicating important contributions from local RNA structure and sequence context. Consistent with previous work, nucleotides flanking the edited adenosine also influenced activity, with favorable editing observed most frequently at 5′-CAN-3′ motifs as shown in EGFP targets^15,32^. These features provide practical design rules for future optimization of Rptr-based editors.

Engineering the Cas9 scaffold further expanded the platform. Deleting the PI domain abolished activity, whereas mutating two inactivating mutations was tolerated, indicating that structural integrity of this region is more important than PAM recognition itself for RNA editing. Deletion of the HNH domain from CjeCas9 remained compatible with editing, consistent with previous structural studies that used an HNH-deleted CjeCas9 construct while preserving the overall protein architecture^22^,.

When benchmarked against REPAIR-v1, RETREAT-Sp1 achieved comparable on-target editing at ACTB while producing substantially fewer transcriptome-wide off-target events, indicating a favorable balance between activity and specificity. Both systems use ADAR2dd carrying the activity-enhancing E488Q mutation. Although dPspCas13b–ADAR2dd was expressed at approximately twice the level of dSpyCas9–ADAR2dd, this difference may only partly account for the greater off-target activity of REPAIR-v1. Differences in the targeting scaffold and fusion architecture may also influence ADAR2dd access to non-target RNAs. Additionally, previous work showed that adding the T375G mutation to ADAR2dd(E488Q) generated the higher-specificity REPAIR-v2 editor while retaining efficient on-target editing^15^. Whether this balance between activity and specificity can be further improved through optimized Rptr design, higher-specificity deaminases, or expression tuning remains an important question.

Beyond single-base editing, RETREAT also enhanced both 3′ and 5′ RNA trans-splicing, demonstrating that Rptr-guided Cas9 recruitment can support larger-scale transcript rewriting. The dependence on guide placement and binding-domain architecture indicates that spatial organization of the editing complex is a key determinant of splicing outcomes, consistent with previous dCas13-assisted trans-splicing studies^25^. Future work should test this strategy at endogenous disease-relevant loci and explore additional Cas9-linked effectors for cleavage, other modifications or RNA modulation. Collectively, our findings establish reprogrammed tracrRNAs as a versatile route to programmable RNA manipulation.

## ACKNOWLEDGMENTS

We thank the Genome Analytics Core Unit facility at Helmholtz Centre for Infection Research for providing the high-throughput sequencing services. The work was supported by funding through the European Research Council (865973 to C.L.B.). C.P. acknowledges funding from the Research Council of Lithuania under the Programme University Excellence Initiatives of the Ministry of Education, Science and Sports of the Republic of Lithuania (Measure No. 12-001-01-01-01, Improving the Research and Study Environment), project No: S-A-UEI-23-10.

## CONFLICTS OF INTEREST

A.A. and C.L.B. have filed a related patent application. C.L.B. is a co-founder of Leopard Biosciences and a co-founder and Scientific Advisory Board member of Locus Biosciences. The other authors declare no competing interests.

## DATA AVAILABILITY

Raw sequencing reads have been deposited in the NCBI Sequence Read Archive (SRA) database under accession number PRJNA1366071 and will be made publicly available upon publication of the peer-reviewed manuscript. Source data underlying all plotted results are provided with this preprint.

## CODE AVAILABILITY

All scripts used for the analysis of processed next-generation sequencing data were deposited to GitHub (https://github.com/beisellab/RETREAT) and will be made publicly available upon publication of the peer-reviewed manuscript.

## AUTHOR CONTRIBUTIONS

Conceptualization: A.Q.A. and C.L.B.; Methodology: A.Q.A.; Formal Analysis: A.Q.A.; Investigation and experimentation: A.Q.A.; Technical assistance: R. L.; Writing – Original Draft: A.Q.A., C.P., and C.L.B.; Writing – Review and Editing: A.Q.A., C.P., and C.L.B.; Visualization: A.Q.A. and C.P.; Supervision: C.L.B.; Funding Acquisition: C.L.B.

## METHODS

### Mammalian cell culture

HEK293T cells (ATCC CRL-11268) (**Table S1**) were cultured in DMEM (Life Technologies) supplemented with 10% (v/v) fetal bovine serum (Corning or BANF Biotrend), 1× penicillin–streptomycin (Life Technologies), and 2□mM L-glutamine. Cells were maintained at 37□°C with 5% CO□ in humidified incubators and passaged at 70–90% confluency using 0.05% Trypsin–EDTA.

### Design and cloning of plasmids for RNA editing

Primers were purchased from Integrated DNA Technologies (IDT). Plasmids and Benchling links are listed in **Table S2**, and Rptr and gRNA sequences are listed in **Table S4**. Constructs were generated using NEBuilder HiFi DNA Assembly Master Mix (NEB, E2621) or Q5 Site-Directed Mutagenesis Kit (NEB, E0554S) and transformed into chemically competent *E. coli* DH5α or TOP10.

The REPAIR-v1 plasmid pC0048 encoding dPspCas13b–ADAR2dd(E488Q) (Addgene, 103864)^15^ was used as a control and as the backbone for dSpyCas9–ADAR2dd constructs. Humanized SpyCas9 from pX330 (Addgene, 42230)^33^ was rendered catalytically inactive by introducing D10A and H840A mutations and used to replace dPspCas13b. Humanized CjeCas9 from pCAG-hCjeCas9-NLS-3×FLAG (Addgene, 133789)^34^ was inactivated by introducing D8A and H559A mutations, and domain-minimized variants were generated by targeted deletion and linker insertion.

For reporter assays, the U6-gRNA cassette from pC0043 (Addgene, 103854)^15^ was inserted into the mCherry-P2A-EGFP(W58X) (Addgene, 172229)^35^ reporter. For dSpyCas9 assays, the gRNA cassette was replaced with the SpyCas9 tracrRNA scaffold and the indicated Rptr variants. For endogenous RNA editing, the U6-Rptr cassette and dCas9–ADAR2dd-P2A-EGFP were combined in a single plasmid.

### Transient transfection and flow cytometry for EGFP rescue

Cells were seeded in 48-well plates 24 h before transfection. At 70–80% confluency, 0.064 pmol each of the Rptr/gRNA-EGFP reporter and dCas–ADAR2dd(E488Q) plasmids were transfected per well using Lipofectamine 3000, according to the manufacturer’s instructions.

After 48 h, cells were dissociated using Trypsin–EDTA and analyzed on a NovoCyte Quanteon cytometer using mCherry (615□nm) and EGFP (530□nm) channels. Debris and doublets were excluded, transfected cells were identified by mCherry expression, and editing was quantified as the percentage of EGFP-positive cells within the mCherry-positive population. Approximately 70,000–100,000 events were acquired per sample. Data was analyzed using NovoExpress and plotted in GraphPad Prism.

### RNA extraction and Sanger sequencing

RNA was isolated using TRI reagent (Zymo, R2050) and the Direct-zol RNA MiniPrep Kit (Zymo, R2052) and eluted in nuclease-free water. Concentrations were measured using NanoDrop 2000 (Thermo Scientific). For plasmid-encoded transcripts, ∼500□ng total RNA and for endogenous transcripts ∼1□µg total RNA was reverse transcribed using SuperScript IV™ (Invitrogen, 18090050) and gene-specific primers (**Table S3**). cDNA was amplified for 35 cycles, purified using DNA Clean & Concentrator-5 (Zymo, D4004), and analyzed by Sanger sequencing. Editing efficiencies were quantified using EditR^18^.

### Targeted RNA editing of endogenous transcripts

Cells were seeded in 24-well plates and transfected at 70–80% confluency with 0.5 µg CMV-dCas9/13–ADAR2dd-P2A–EGFP plasmid and the corresponding U6-Rptr/gRNA plasmid using Lipofectamine 3000. After 48 h, cells were harvested and total RNA was isolated, reverse-transcribed, and processed as described above.

Target regions were amplified by two-step PCR using KAPA HiFi HotStart ReadyMix, with 15 cycles using gene-specific primers containing partial Illumina adapters, followed by 15 cycles to add indexed adapters (**Table S3**). Libraries were purified, pooled equimolarly, and sequenced on an Illumina NovaSeq to generate at least 1 million 150-bp paired-end reads per sample.

Reads were adapter- and quality-trimmed with Trimmomatic v0.36^36^, aligned to gene-specific amplicon references using BWA-MEM v0.7.15^37^, and processed with Samtools v1.21^38^. Base counts were obtained using samtools mpileup with minimum mapping and base quality scores of 20. A custom Python script quantified A and G reads at each reference adenosine and calculated A-to-G editing as G/(A+G).

### Whole-transcriptome sequencing for specificity analysis

Total RNA was isolated 48 h after transfection using Direct-zol RNA MiniPrep Kits with on-column DNase I treatment. RNA quality was assessed by NanoDrop, Qubit, and Fragment Analyzer, and samples with RQN >8 were used for library preparation. Poly(A)+ RNA was enriched from 500□ng total RNA using the NEBNext Poly(A) mRNA Magnetic Isolation Module (NEB, E3370), and directional libraries were prepared using the NEBNext Ultra II kit (NEB, E7760). Libraries were sequenced on an Illumina NovaSeq X to generate approximately 40 million 150-bp paired-end reads per sample.

### Transcriptome-wide RNA editing analysis

After demultiplexing, reads were trimmed with fastp v0.22.0^39^ and aligned in two-pass mode to GRCh38 supplemented with the Cas–ADAR2dd sequence using STAR v2.7.10b^40^. Alignments were processed with Samtools v1.21^38^ and downsampled to 35 million properly paired reads per sample.

RNA-editing sites were identified with REDItools2^41^ using minimum mapping and base quality scores of 30 and 20, respectively, a minimum depth of 10, and at least three edited reads. Only A-to-G and T-to-C substitutions were retained. Sites overlapping common dbSNP variants, RepeatMasker-annotated regions, or Cas–ADAR2dd sequences were excluded. High-confidence sites were defined using a 95% Wilson confidence interval and retained only if detected in all biological replicates. Site overlaps were visualized with DeepVenn^42^, and transcript abundance was quantified with Salmon v1.10.3^43^ and reported as TPM.

### Construct design for RNA trans-splicing

Primers were purchased from Integrated DNA Technologies. Plasmids and Benchling links are listed in **Table S2**, and trans-splicing sequences are listed in **Table S5**. The 1×NLS–dSpyCas9–2×NLS construct was generated from CBS-5067 by replacing the NES–ADAR2dd cassette with an additional NLS.

For 3′ trans-splicing, the target plasmid contained MMP9 intron 1 and exon 2 downstream of a CMV-driven mCherry–P2A–5′-EGFP cassette. Primer-binding sites were introduced into MMP9 exon 2 to enable amplification of cis- and trans-spliced products. The donor plasmid encoded a 20–50-nt binding domain, a 56-nt 3′ hemi-intron, and 3′-EGFP–IRES–BFP, together with a U6-driven Rptr cassette.

For 5′ trans-splicing, the target plasmid contained MMP9 exon 1, intron 1, and 3′-EGFP–IRES–BFP. The donor encoded mCherry–5′-EGFP, a 25-nt 5′ hemi-intron, and a binding domain. In design 1, the Rptr was expressed separately from a U6 promoter, whereas in design 2, it was fused directly to the donor transcript.

### Split-EGFP reporter assay for RNA trans-splicing

Cells were seeded in 24-well plates 24 h before transfection. At 70–80% confluency, 0.064 pmol each of the Rptr/gRNA-EGFP reporter and dCas–ADAR2dd(E488Q) plasmids were transfected per well using Lipofectamine 3000, according to the manufacturer’s instructions.. DNA and reagent mixtures were prepared according to the manufacturer’s protocol. When dSpyCas9 was omitted, an equimolar amount of dCjeCas9-NES was included as a size-matched control.

Cells were incubated for 48□h, dissociated with Trypsin, neutralized with PBS containing 10% FBS, and collected for flow cytometry and RNA extraction.

### Library preparation for targeted sequencing of trans-splicing assays

RNA extraction, cDNA synthesis, and two-step amplicon library preparation were performed as described for endogenous targeted editing. The libraries were pooled in equimolar amounts and at least 1 million 150□bp paired-end reads were generated for each sample, using Illumina NovaSeq X.

### Analysis of RNA trans-splicing products

Paired-end reads were aligned to 150-bp cis- and trans-spliced reference amplicons using Bowtie2 v2.5.4^44^ in very-sensitive end-to-end mode. Only uniquely mapped reads with MAPQ ≥30, ≥95% identity, and sufficient junction-spanning sequence were retained. A minimum product-specific overhang of 20 bp was required for 3′ trans-splicing and 25–30 bp for 5′ trans-splicing. Soft-clipped and multi-mapping reads were excluded. Reads were counted with featureCounts v2.1.1^45^, and trans-splicing efficiency was calculated as trans-spliced reads divided by the sum of trans- and cis-spliced reads.

