## Supplementary Information for "RNA editing and trans-splicing with reprogrammed tracrRNAs"

**Reprogrammed tracrRNA enables Cas9-mediated RNA editing and trans-splicing**

SUPPLEMENTARY FIGURES	2

SUPPLEMENTARY TABLES	13

SOURCE DATA	13

SUPPLEMENTARY DATASET	13

### SUPPLEMENTARY FIGURES

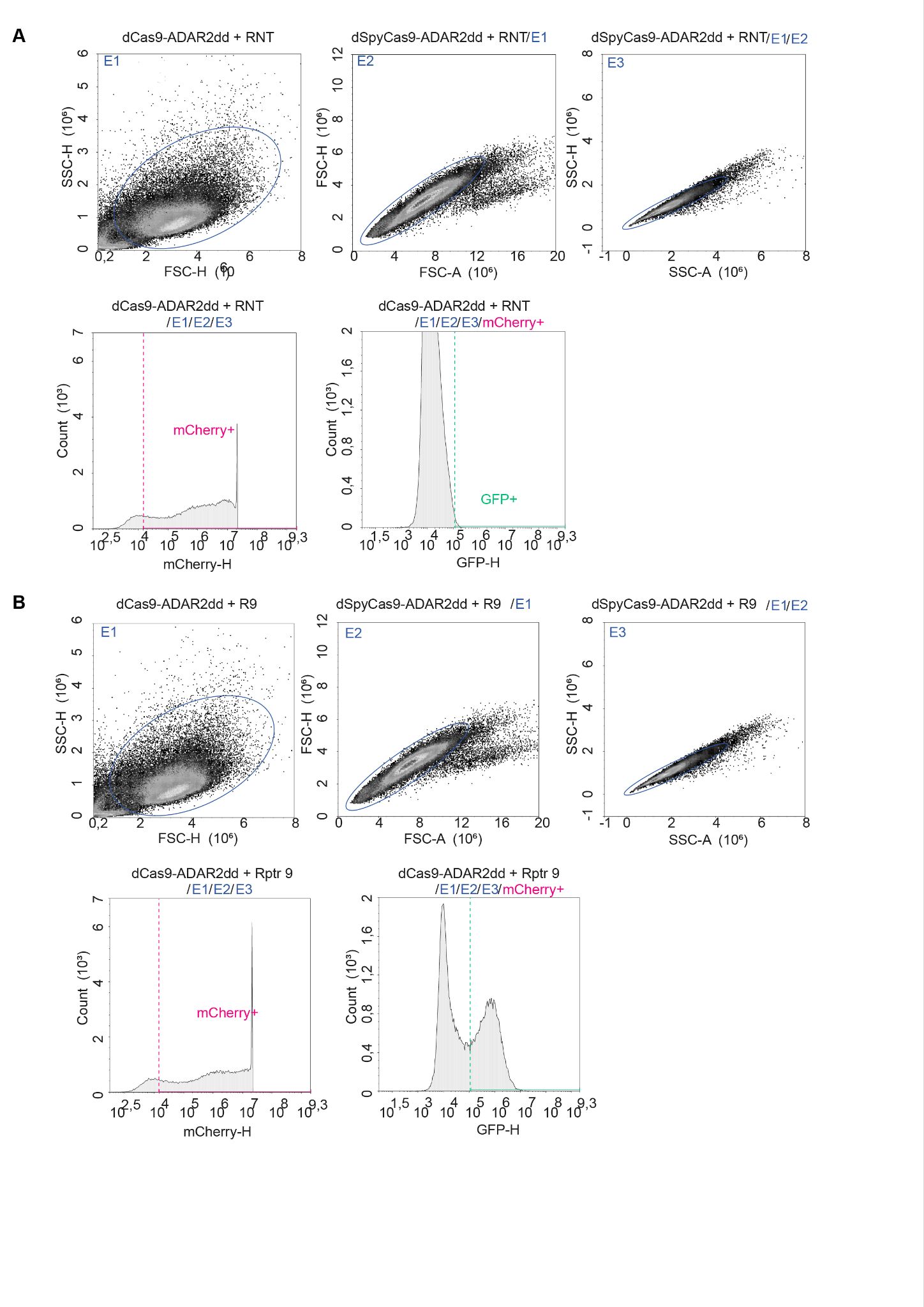

#### Fig. S1 | Gating strategy for flow cytometry–based EGFP(W58X) reporter analysis.

(**A**) Representative flow cytometry plots showing the gating strategy used to quantify EGFP restoration. Live cells were first gated, followed by two sequential singlet gates. Transfected cells were identified as mCherry-positive (mCherry+), and editing was measured as the fraction of GFP-positive (GFP+) cells within the mCherry+ population. Fluorescence gates were defined using untransfected, mCherry-only, and mCherry + EGFP controls (not shown). Data shown are from cells transfected with the W58X EGFP reporter, dSpyCas9–ADAR2dd and a non-targeting Rptr (RNT). (**B**) Representative plots showing the same gating workflow applied to cells transfected with the W58X EGFP reporter, dSpyCas9–ADAR2dd and the targeting Rptr 9 (R9).

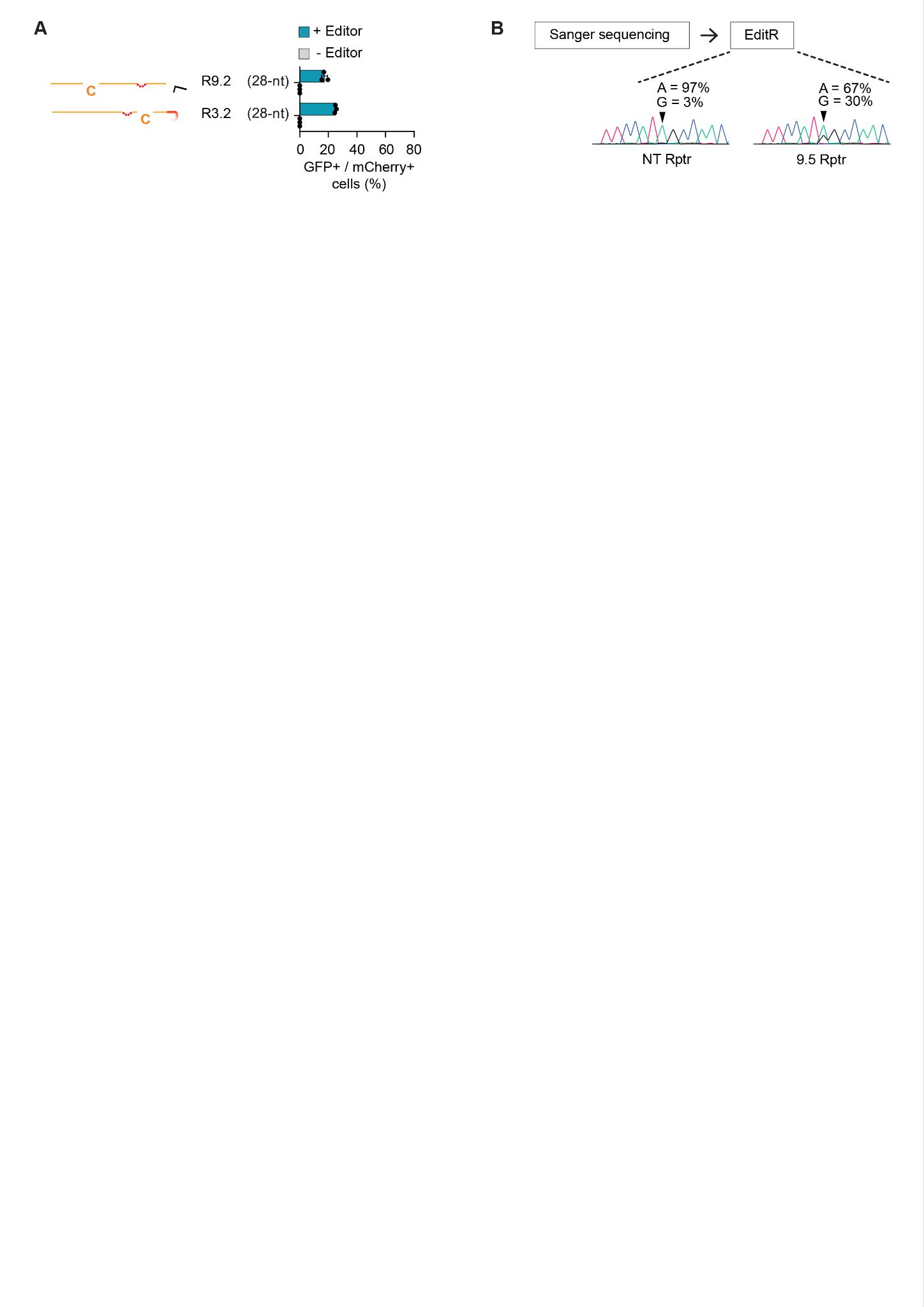

#### Fig. S2 | Editing outcomes of shortened Rptr designs and representative Sanger traces.

(**A**) Editing efficiencies of Rptr 3 and Rptr 9 variants with programmable regions shortened by 10 nt. Bars show RNA editing with (teal) or without (gray) the editor. Data represent mean ± SD (n = 3), with individual replicates shown as dots. (**B**) Representative Sanger sequencing chromatogram illustrating the double peak characteristic of A-to-I(G) editing after EditR analysis.

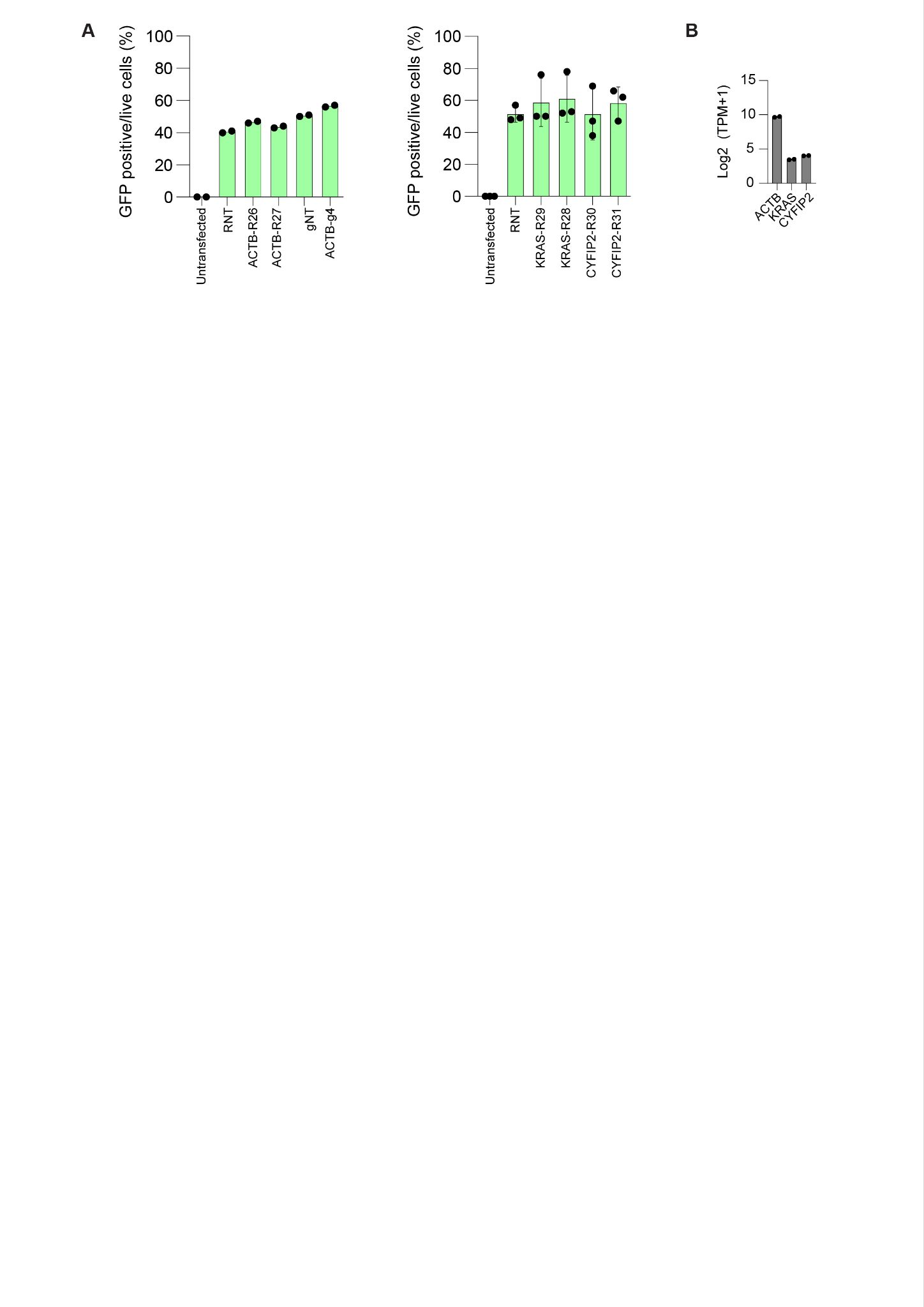

#### Fig. S3 | Transfection efficiency, normalized on-target editing, and endogenous expression levels.

(**A**) Transfection efficiency of editor–Rptr plasmids, measured as the percentage of GFP-positive cells among live cells. GFP is expressed downstream of the editor via a P2A sequence. Data are shown for ACTB targeting using RETREAT-Sv1 and REPAIR-v1, and for KRAS and CYFIP2 targeting using RETREAT-Sv1. (**B**) Baseline expression levels of ACTB, KRAS and CYFIP2 in untransfected HEK293T cells, shown as log2(TPM + 1).

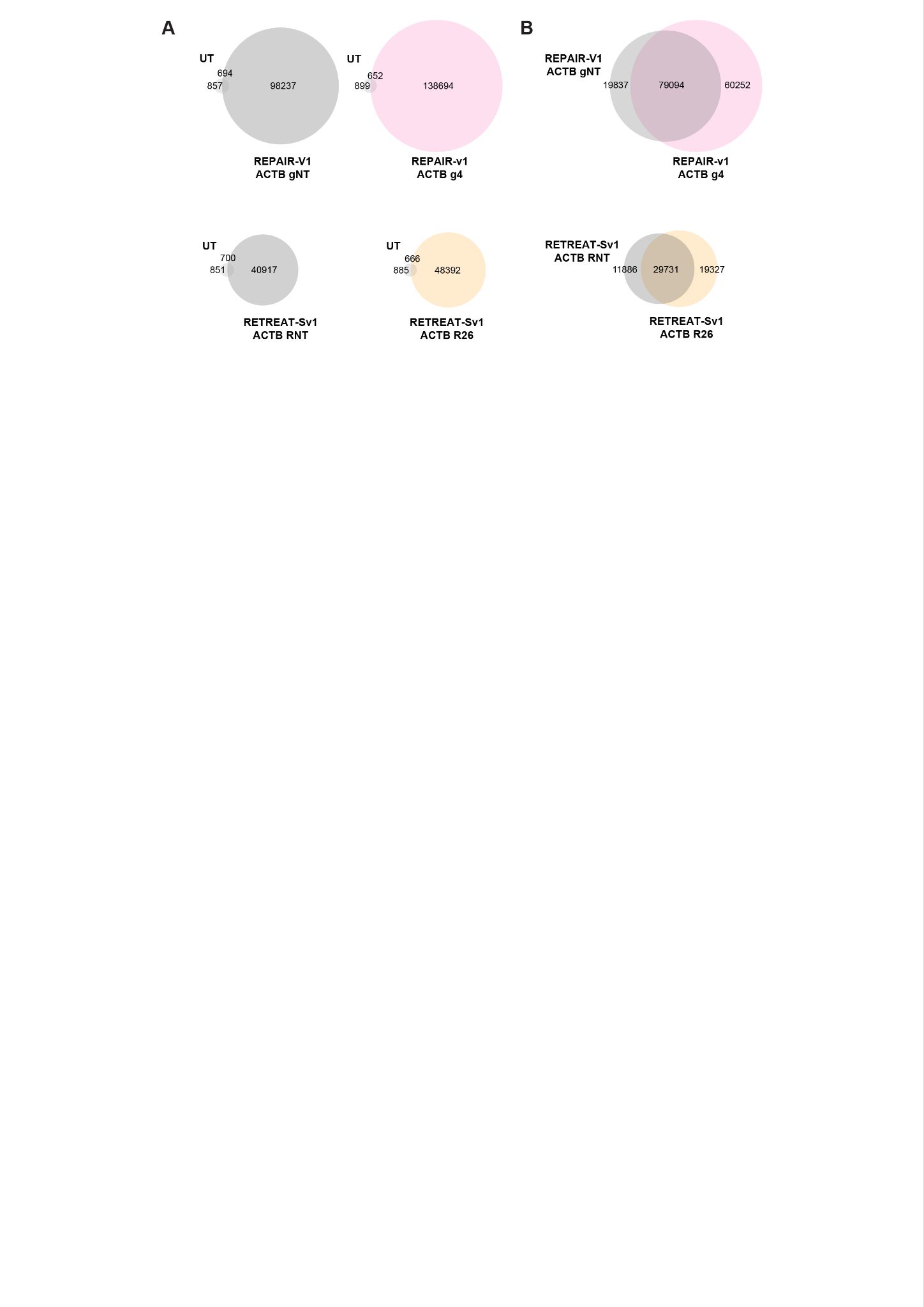

#### Fig. S4 | Overlap of transcriptome-wide off-target edits across conditions.

(A) Venn diagrams showing shared A-to-I(G) editing sites between untransfected HEK293T cells and each editor condition (REPAIR-v1 or RETREAT-Sv1) with either an ACTB-targeting guide or a non-targeting (NT) guide. (B) Venn diagrams comparing off-target edits between NT and targeting guides within each editor: REPAIR-v1 (NT vs g4) and RETREAT-Sv1 (NT vs R26).

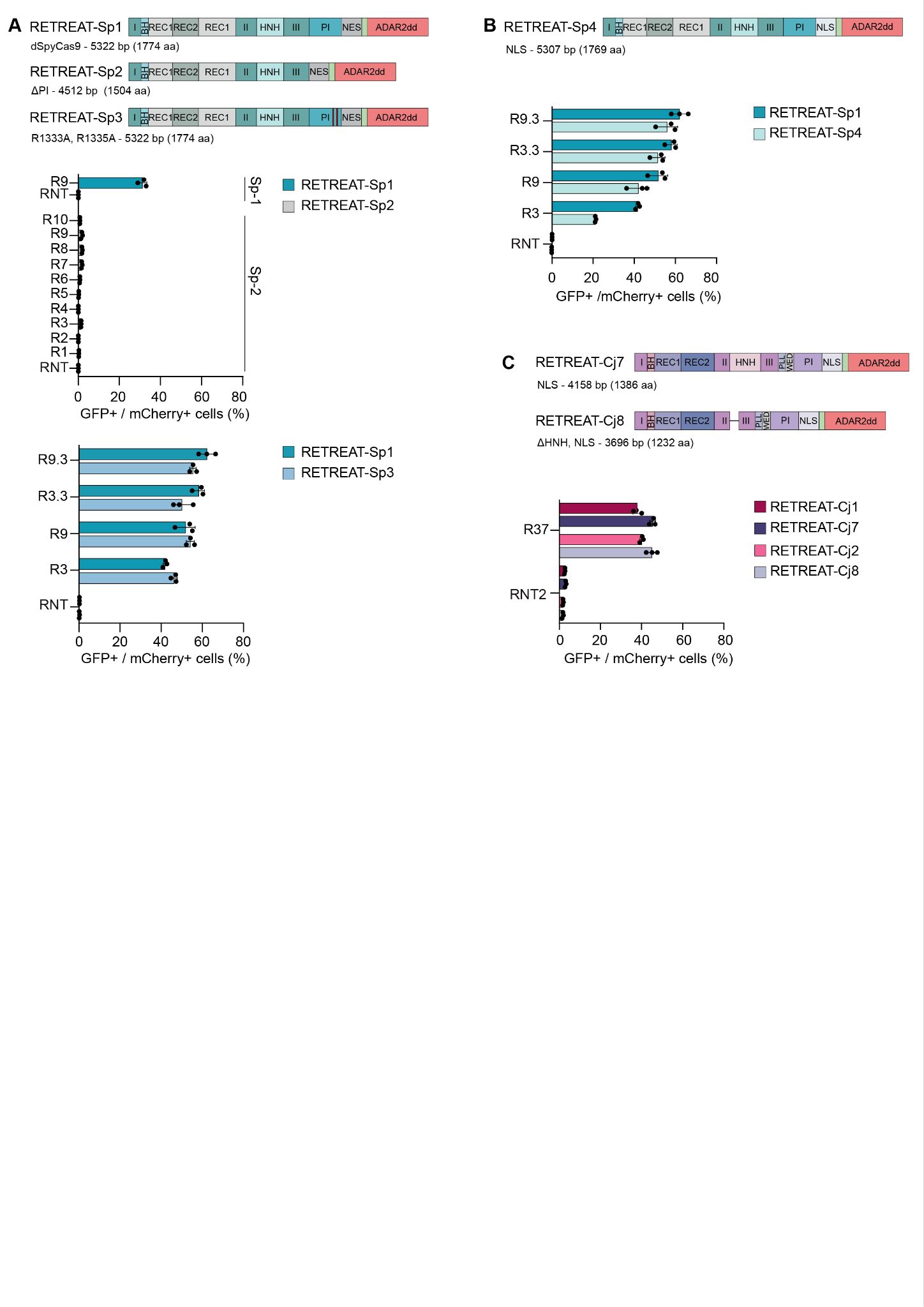

**Fig. S5 | Effects of Cas9 engineering and subcellular localization on RETREAT-mediated RNA editing.**

(A) RNA editing with SpyCas9 variants carrying PI-domain deletions or PAM-interacting mutations.

(B) Effect of replacing the nuclear export signal (NES) with a nuclear localization signal (NLS) in SpyCas9-based RETREAT. The NES-to-NLS substitution reduced editing from 41.9% to 21.3% with Rptr 3, from 58.3% to 51.7% with Rptr 3.3, from 51.9% to 42.3% with Rptr 9, and from 62.3% to 56.2% with Rptr 9.3.

(C) Effect of replacing the NES with an NLS in CjeCas9-based RETREAT. Nuclear localization increased editing from 37.9% to 45.4% for full-length dCjeCas9 and from 40.1% to 45.1% for the ΔHNH variant. Together, these results indicate that the effect of nuclear localization on RETREAT activity depends on the Cas9 ortholog.

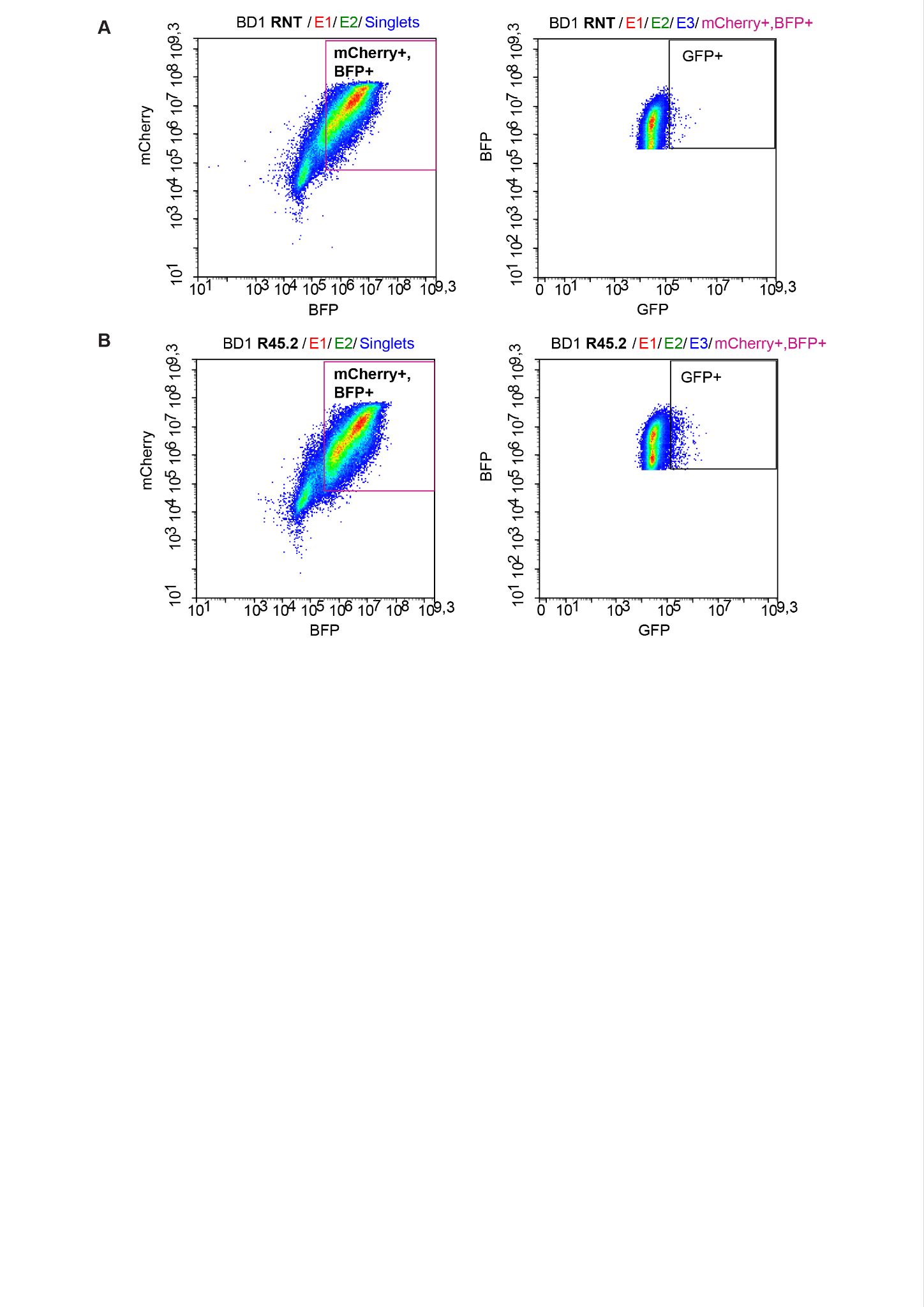

#### Fig. S6 | Gating strategy for 3′ trans-splicing analysis.

(**A**) Representative gating for BD1 with a non-targeting Rptr. Live cells and singlets were first selected as in Fig. S1. Transfected cells were identified using a two-dimensional gate on the mCherry (y-axis) and BFP (x-axis) fluorescence plot to isolate the double-positive population. Trans-splicing efficiency was quantified as the percentage of GFP-positive cells within this mCherry+/BFP+ gate. (**B**) Representative gating for BD1 with the targeting Rptr 45.2, showing GFP-positive cells within the mCherry+/BFP+ population.

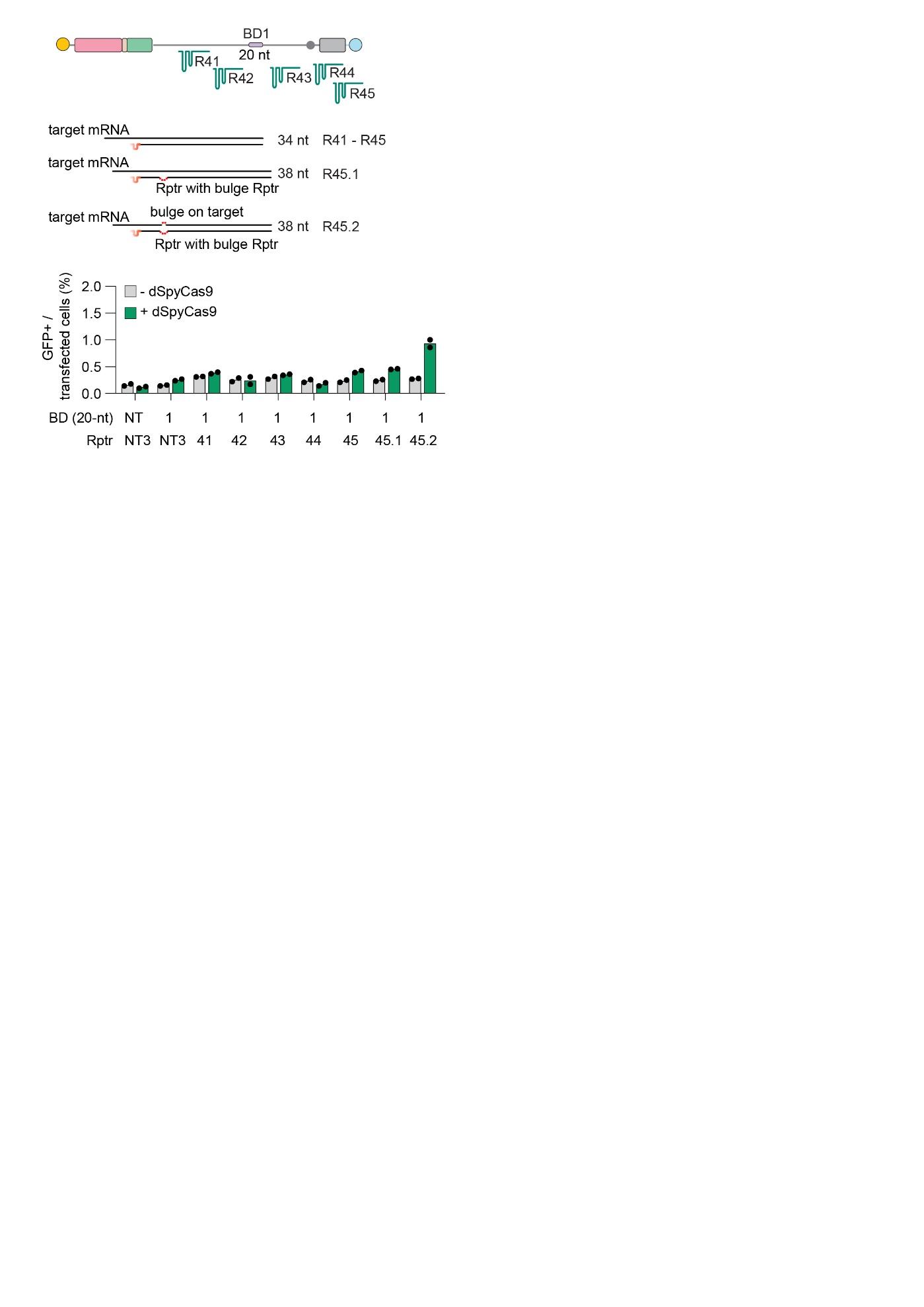

#### Fig. S7 | Initial optimization of Rptr design and dCas9 recruitment for BD1-mediated 3′ RNA trans-splicing.

BD1-mediated 3′ RNA trans-splicing using Rptrs 41–45, 45.1, and 45.2. Controls included a non-targeting BD, a non-targeting Rptr, and omission of dCas9. Trans-splicing efficiency was quantified by flow cytometry as the percentage of GFP-positive cells within the mCherry- and BFP-double-positive population.

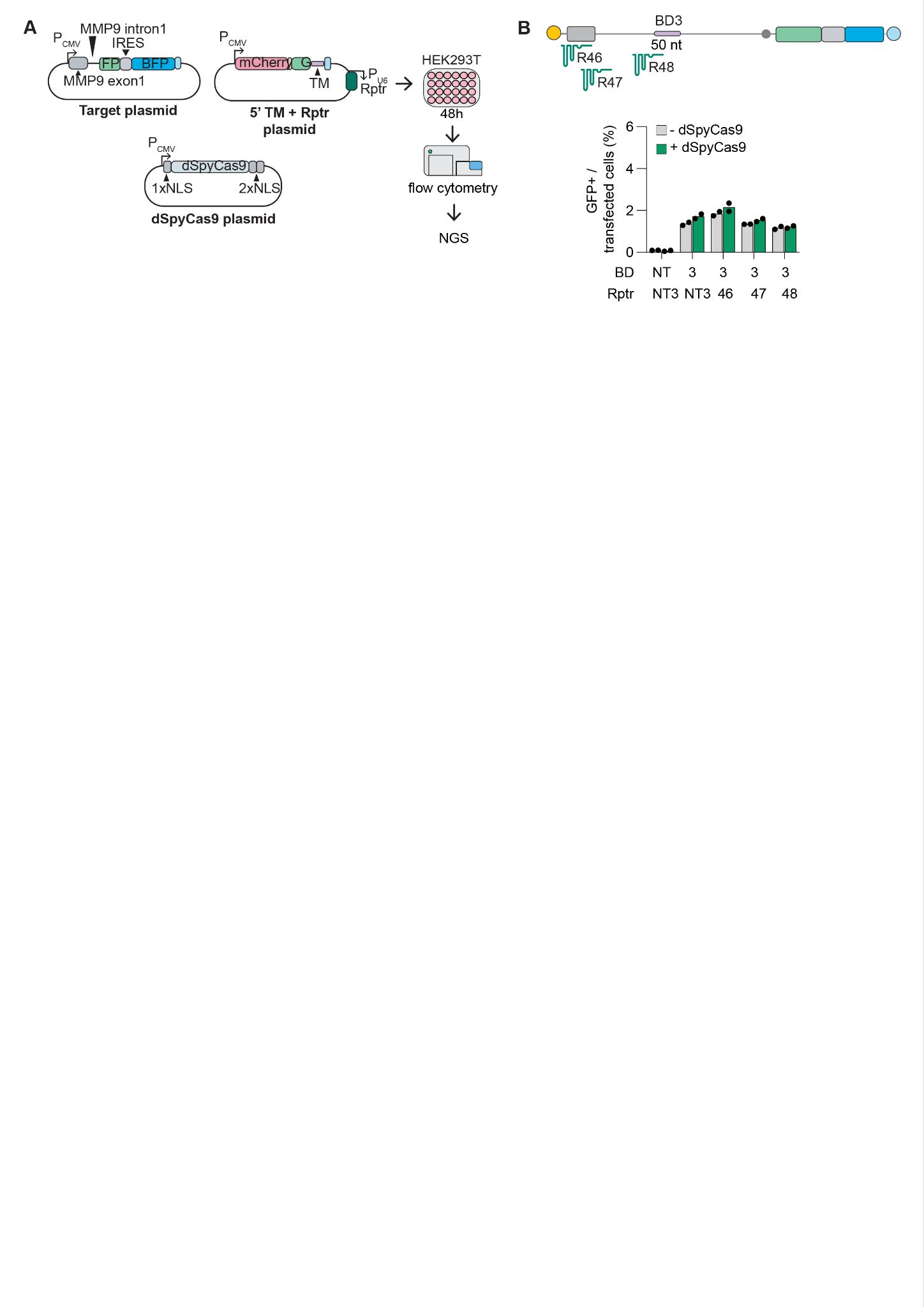

#### Fig. S8 | Initial design and testing of RETREAT for 5′ RNA trans-splicing.

(**A**) Schematic of the plasmids and workflow used for 5′ RNA trans-splicing. HEK293T cells were transfected with plasmids encoding the reporter acceptor, the donor containing the Rptr, 5′ cargo, and trans-splicing molecule (TM), and dCas9-NLS, followed by flow cytometry and targeted NGS.
(**B**) BD3-mediated 5′ RNA trans-splicing using Rptrs 46–48. Controls included a non-targeting Rptr and omission of dCas9. Trans-splicing efficiency was quantified by flow cytometry as the percentage of GFP-positive cells within the mCherry- and BFP-double-positive population.

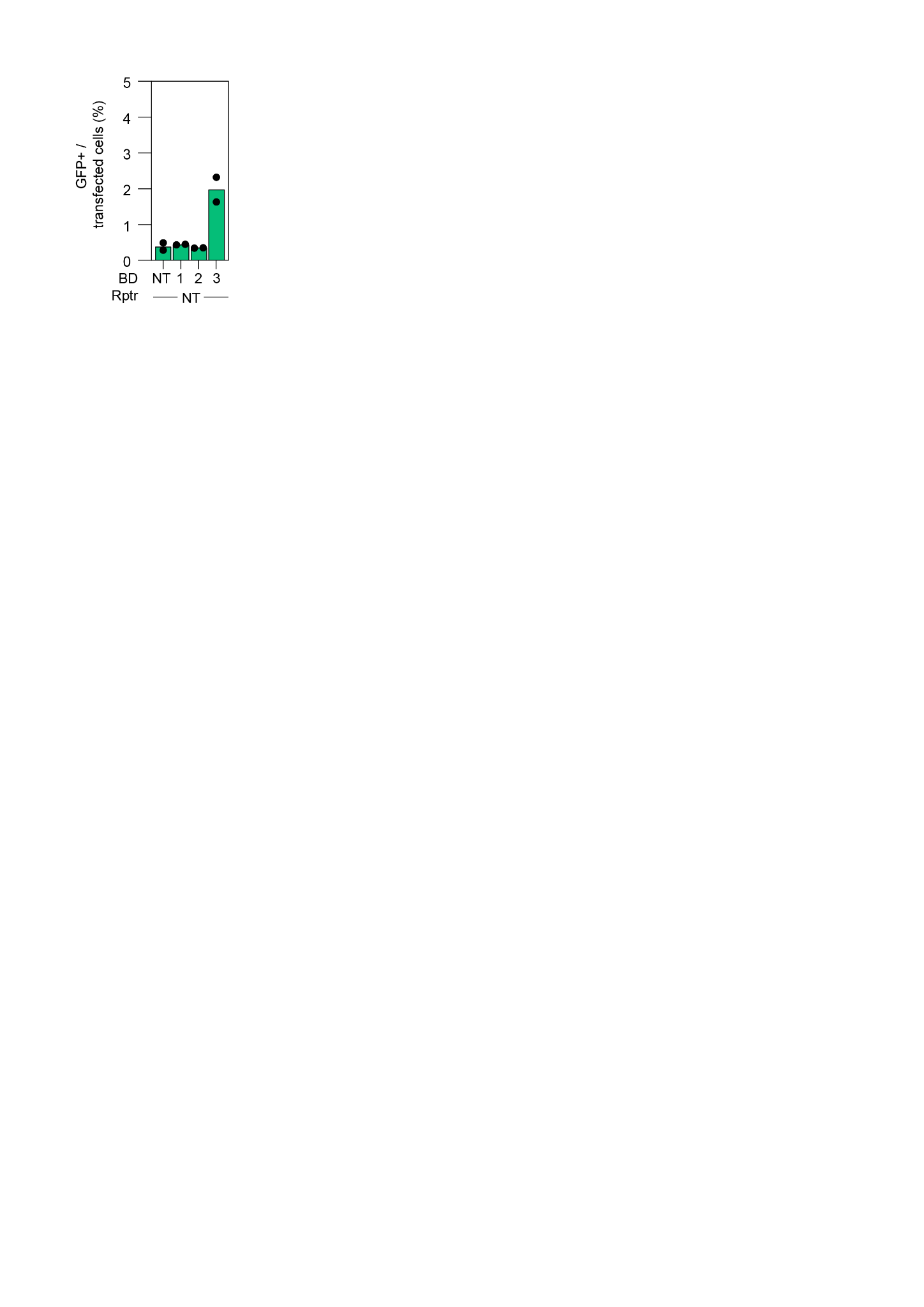

#### Fig. S9 | Baseline 5′ trans-splicing efficiencies of binding domain variants.

GFP-positive cells among the transfected population were measured to assess basal 5′ trans-splicing activity for non-targeting BD, BD1, BD2 and BD3 in the presence of dCas9. Bars show the percentage of GFP-positive cells within the mCherry+/BFP+ double-positive gate, indicating trans-splicing efficiency. BD3 exhibited the highest basal activity, whereas BD1, BD2 and the non-targeting BD showed similarly low levels (<0.5%). Bar plots represent mean (n = 2), with individual replicates shown as dots.

### SUPPLEMENTARY TABLES

**Table S1.** List of bacterial strains and mammalian cell lines used in this study.

**Table S2.** List of plasmids used in this study.

**Table S3.** List of important oligonucleotides used in this study.

**Table S4.** List of Rptr/gRNA spacer sequences used in this study.

**Table S5.** List of important sequences for trans-splicing experiments including binding domain (BD) and hemi-intron.

### SOURCE DATA

All source data underlying the figures presented in the Results section (e.g., flow-cytometry–based editing efficiencies, EditR outputs, and on-target NGS analyses).

### SUPPLEMENTARY DATASET

**Supplementary Dataset 1** – Transfection efficiencies corresponding to Fig. 1, Fig. 2, Fig. 4, Fig. S2, and Fig. S5.

**Supplementary Dataset 2** – Transcriptome-wide editing events detected in UT, RETREAT-Sv1 (R26 and RNT), and REPAIR-v1 (g4 and gNT) conditions.

**Supplementary Dataset 3** – Merged transcript-level TPM matrix derived from Salmon quantifications across all experimental conditions.
